# Maternal cannabis vapor exposure causes long-term alterations in emotional reactivity, social behavior, and behavioral flexibility in offspring

**DOI:** 10.1101/2020.03.12.989210

**Authors:** Halle V. Weimar, Hayden R. Wright, Collin R. Warrick, Amanda M. Brown, Janelle M. Lugo, Timothy G. Freels, Ryan J. McLaughlin

## Abstract

The use of cannabis during pregnancy is a growing public health concern. As more states implement legislation permitting recreational cannabis use, there is an urgent need to better understand its impact on fetal neurodevelopment and its long-term effects in exposed offspring. Studies examining effects of prenatal cannabis exposure typically employ injections of synthetic cannabinoids or isolated cannabis constituents that may not accurately model cannabis use in human populations. To address this limitation, we have developed a novel e-cigarette technology-based system to deliver vaporized cannabis extracts to pregnant Long Evans rats. We used this model to determine effects of prenatal cannabis exposure on emotional, social, and cognitive endpoints of male and female offspring during early development and into adulthood. Dams were exposed to cannabis vapor (CAN_THC_: 400 mg/ml), vehicle vapor (VEH), or no vapor (AIR) twice daily during mating and gestation. Offspring exposed to CAN_THC_ and VEH showed reduced weight gain relative to AIR offspring prior to weaning. CAN_THC_ offspring made more isolation-induced ultrasonic vocalizations (USVs) on postnatal day 6 (P6) relative to VEH-exposed offspring, which is indicative of increased emotional reactivity. Male CAN_THC_ offspring engaged in fewer social investigation behaviors than VEH-exposed male offspring during a social play test on P26. In adulthood, CAN_THC_-exposed offspring spent less time exploring the open arms of the elevated plus maze and exhibited dose-dependent deficits in behavioral flexibility in an attentional set-shifting task relative to AIR controls. These data collectively indicate that prenatal cannabis exposure causes enduring effects on the behavioral profile of offspring.

## 1. INTRODUCTION

The prevalence of cannabis use during pregnancy in the United States has more than doubled over the past two decades (Volkow et al., 2019). Nationwide, 7% of pregnant women report using cannabis, which makes it the most commonly used illicit drug during pregnancy (Volkow et al., 2019). Approximately 70% of women believe that there is little or no risk in using cannabis once or twice a week during pregnancy (Ko et al., 2015). Additionally, cannabis use among pregnant women is becoming more prevalent, in part due to its antiemetic properties (Roberson et al., 2014; Westfall et al., 2006) and endorsements from cannabis dispensaries (Dickson et al., 2018). As the perceived harm and social stigma associated with cannabis use continue to decline (Wen et al., 2019), there is significant concern that the use of cannabis among pregnant women will continue to rise in the coming years. This is particularly troubling since the long-term ramifications of prenatal cannabis exposure remain largely unknown.

The main psychoactive component in cannabis, Δ^9^-tetrahydrocannabinol (THC), is known to cross the placenta (Bailey et al., 1987), and can interfere with endogenous cannabinoid signaling, which is critically involved in several neurodevelopmental processes (see Scheyer et al., 2019b for review). Our understanding of the developmental effects of prenatal cannabis exposure has largely been informed by human longitudinal studies of cannabis-exposed offspring (Fried and Watkinson, 2001; Fried et al., 1998, 2003; Richardson et al., 2002; Smith et al., 2016). These studies have revealed associations between prenatal cannabis exposure and deficits in cognition (Fried et al., 1998), executive functioning (Smith et al., 2016), and working memory (Fried and Watkinson, 1990). Cross-sectional retrospective studies have also linked prenatal cannabis exposure to impaired memory, impulse control, problem solving, and quantitative reasoning (Sharapova et al., 2018), as well as alterations in emotional reactivity and increased symptoms of affective disorders (De Moraes Barros et al., 2006; Fried and Makin, 1987; Leech et al., 2006; Lester and Dreher, 1989). However, the interpretation of the human literature is complicated by confounding factors, including underreporting of use (Metz et al., 2017) and co-use of other drugs (Qato et al., 2020). Thus, very little is known about the enduring effects of prenatal cannabis exposure, independent of these extraneous factors.

Animal models provide an opportunity to explore causal effects of prenatal cannabis exposure without the confounds inherent to human studies. Models of prenatal cannabis exposure in rodents have supported alterations or deficits across cognitive, emotional, social, and motor domains in offspring (Antonelli et al., 2005; Campolongo et al., 2011; Manduca et al., 2020; Navarro et al., 1994, 1996; Newsom and Kelly, 2008; Trezza et al., 2008). Moreover, lasting changes in epigenetic regulation (DiNieri et al., 2011; Spano et al., 2007), synaptic plasticity (Bara et al., 2018; Scheyer et al., 2019a), and dopamine neuron ontogeny/signaling (Bonnin et al., 1994, 1995, 1996; DiNieri et al., 2011; Frau et al., 2019; Rodríguez de Fonseca et al., 1990, 1991, 1992; Wang et al., 2004) have been observed in rodents that have been perinatally exposed to isolated THC or the CB1 receptor agonist WIN 55,212-2. However, intraperitoneal injections of synthetic cannabinoids may not fully recapitulate the effects of whole-plant cannabis, which contains a multitude of phytocannabinoids beyond THC that each have unique pharmacological and physiological effects (Morales et al., 2017; Russo et al., 2011; Turner et al., 2017). Moreover, synthetic CB1 receptor agonists are known to recruit different intracellular signaling cascades (Laprairie et al., 2014, 2016) and differences in pharmacokinetics due to the route of administration can dramatically influence the amount of fetal exposure (McLaughlin, 2018). More recently, a cannabis vapor delivery system has been introduced to more closely mimic the intrapulmonary route of administration that is most common among human cannabis users (Freels et al., 2020; Javadi-Paydar et al., 2018; Nguyen et al., 2016). This approach takes advantage of commercial ‘e-cigarette’ technology to deliver vaporized whole-plant cannabis extracts to rodents that results in pharmacologically and behaviorally relevant cannabinoid concentrations in plasma and brain tissue (Freels et al., 2020; Nguyen et al., 2016). Implementation of this more ecologically valid approach could provide translational insight into the long-term effects of developmental cannabis exposure (McLaughlin, 2018).

In the current study, we exposed female Long Evans rats to either whole-plant cannabis extract vapor (CAN_THC_), vehicle vapor (VEH), or no vapor (AIR) twice daily prior to, and during, gestation. To test our hypothesis that prenatal cannabis exposure causes enduring effects on the behavioral profile of offspring, we conducted tests of emotional reactivity, social and novel environment-induced anxiety-like behavior, and behavioral flexibility at neonatal, juvenile, and adult timepoints, respectively. Our results indicate that prenatal cannabis exposure increases isolation-induced ultrasonic vocalizations in neonates, negatively impacts social play and investigation at the juvenile stage, and alters anxiety-like behavior induced by novel, anxiogenic environments in adulthood. Moreover, we found that offspring prenatally exposed to cannabis of both sexes exhibited impaired performance in an automated attentional set-shifting task designed to assess behavioral flexibility in adulthood. Thus, maternal cannabis vapor exposure causes age-dependent effects in exposed offspring that extend into adulthood.

## 2 MATERIALS AND METHODS

### 2.1. Animals

Male and female Long Evans rats (Simonsen Laboratories, Gilroy, CA; 60-70 day old) were same-sex, pair-housed in a humidity-controlled animal room on a 12:12 reverse light cycle (7:00 lights off, 19:00 lights on) with food and water available *ad libitium*. One week after arrival, all females were randomly assigned to CAN_THC_, VEH, or AIR groups and then paired with a male in a Double Decker cage (Tecniplast, Milan, Italy). All females regardless of group were weighed and handled daily. After one week of mating, males were removed and females were single-housed until the day of birth, which was designated as postnatal day 0 (P0). Pups were cross-fostered between P0 and P2 depending on the availability of litters. Before cross-fostering, each litter was weighed and the number and sex of pups was determined. To distinguish prenatal treatment condition, a small ink dot was tattooed on the right, left, or neither hindpaw with Super Black™ india ink (Speedball, Statesville, NC, USA). Litters were culled and subsequently cross-fostered so that each dam raised 10-12 pups. Ideally, each cross-fostered litter contained pups from all three prenatal treatments; however, in some cases only two conditions were raised by each dam due to differences in the timing of births between litters. In one case, a VEH-exposed dam raised her own biological litter of 15 pups due to a lack of available litters for cross-fostering. Litters remained undisturbed (except for ultrasonic vocalization [USV] tests and cage changes) until weaning on P21 when they were group-housed with same-sex littermates. All procedures were performed in accordance with the guidelines in the National Institute of Health Guide for the Care and Use of Laboratory Animals and were approved by the Washington State University Institutional Animal Care and Use Committee.

### 2.2. Prenatal Drug Delivery

#### 2.2.1. Drugs

The whole-plant cannabis extract (CAN_THC_) was obtained from the National Institute on Drug Abuse (NIDA) Drug Supply Program. According to the certificate of analyses provided upon shipment, the CAN_THC_ extract contained 24.8±0.08% THC, 1.2±0.01% cannabidiol (CBD), and 2.1±0.02% cannabinol (CBN). Extracts were heated to 60°C under constant stirring and dissolved in 80% propylene glycol/20% vegetable glycerol at a concentration of 400 mg/ml based on previous studies (Freels et al., 2020; Nguyen et al., 2016). For studies assessing behavioral flexibility, an additional low CAN_THC_ concentration group (50 mg/ml) was included to examine potential dose-dependent effects. The final estimated concentrations of phytocannabinoids in the 400 mg/ml CAN_THC_ preparation were: 99.2 mg/ml THC, 4.8 mg/ml CBD, and 8.4 mg/ml CBN. The final estimated concentrations of phytocannabinoids in the 50 mg/ml CAN_THC_ preparation were: 12.4 mg/ml THC, 0.6 mg/ml CBD, and 1.1 mg/ml CBN.

#### 2.2.2. Vapor Exposure Regimen

Vapor exposure sessions were conducted using 13.5” × 9.0” × 8.25” (L × W × H) 16.4 L vapor chambers from La Jolla Alcohol Research Inc. (La Jolla, CA) with vaporizer boxes controlled by MED Associates IV software (Fairfax, VT). The air intake port pulled air through tubing connected to an air flow meter and tubing connected to a commercial e-cigarette cartridge (first generation: Protank 3 Dual Coil; 2.2 Ω atomizer; KangerTech, Shenzhen, China; second generation: SMOK Tank Baby Beast TFV8 with 0.2Ω M2 atomizer, 40-60 W range) filled with CAN_THC_ or VEH. Starting on the day of pairing, female rats in the two vapor groups began twice-daily vapor exposure sessions, which coincided with the first and last hour of the dark cycle (**Fig. 1**). Vapor was delivered every 2 min for 60 min under continuous rear-port evacuation via a house vacuum system containing in-line Whatman HEPA-Cap filters. For studies using first generation vaporizers and cartridges (i.e., first cohort of adult elevated plus maze [EPM] and first two cohorts of behavioral flexibility studies), vapor was delivered in discrete 10 s puffs. For studies using second generation vaporizers and cartridges (i.e., USVs, social play, juvenile EPM, second adult EPM cohort, and third cohort of behavioral flexibility testing), vapor was delivered in discrete 5 s puffs. Vapor exposure sessions concluded 24-48 hr before the anticipated date of birth.

**Figure 1.**
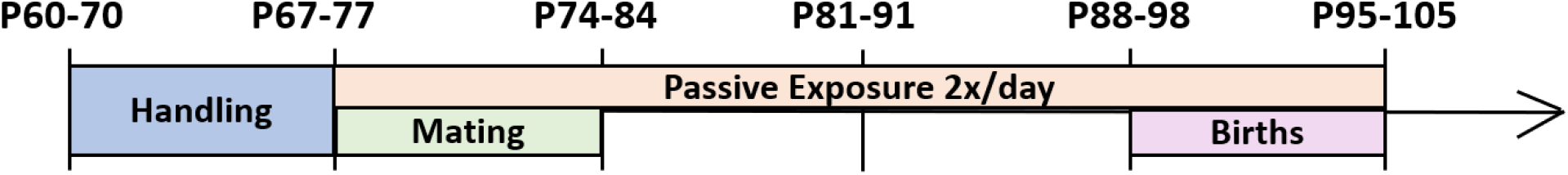
Timeline of vapor exposure paradigm. Schematic timeline of exposure paradigm starting at arrival of rats at facility through cross-fostering on P2.

### 2.3. Behavioral Testing

#### 2.3.1. Ultrasonic Vocalizations

Isolation-induced USVs were recorded from each pup on P6, P10, and P13 as previously described (Hofer et al., 2002). Starting 2 hr into the dark cycle, dams were removed from the home cage and placed in a familiar transfer cage in a separate room. Home cages with litters were transferred to another holding room and placed on a heating pad. Starting after a 10 min habituation period, individual pups were carried to a separate testing room and placed in the center of an empty 17” × 12” × 7.5” (L × W × H) standard rat cage with a 3×3 grid visible underneath. USVs were recorded for 3 min by an UltraSoundGate 116 microphone (Avisoft Bioacoustics, Glienicke/Nordbahn, Germany) suspended 15 cm above the cage floor. Recordings from each session were analyzed offline using DeepSqueak software (Coffey et al., 2019) to determine the number and types of calls. The number of grid crossings per session was determined from the video recordings and scored manually by a researcher who was blinded to treatment condition.

#### 2.3.2. Social Play Behavior

Assessment of social investigation and play behavior was performed on P26 using a protocol modified from Holman et al. (2019). For 2 days prior to testing, juvenile rats were habituated to the play arena for 5 minutes, which was a 24” × 24” × 12” (L × W × H) open field filled with a layer of clean bedding. On the last day of habituation, juvenile rats were single housed overnight to facilitate play behavior during the test phase. On the test day, the fur on both hips of each rat was marked with black markers for identification purposes during later review. Three unfamiliar rats were allowed to interact for 10-min videotaped sessions. Following testing, rats were returned to group housing in their home cage. In rare cases, a rat underwent social play twice on the same day if there were not enough rats to form triads of unfamiliar playmates. In these cases, only the first play session was used for behavioral analysis for this rat. Videos were reviewed by two blinded experimenters who individually recorded time stamps for the following: social investigation behaviors (i.e., anogenital sniffing, body sniffing, social grooming), play behaviors (i.e., dorsal contact, pinning, pouncing, following, evading, wrestling), and asocial behaviors (i.e., digging, self-grooming). The two researchers then reconciled the time stamps together to generate frequency and latency data for each behavioral measure for each rat (see **Table 1** for descriptive statistics). The total number of play, social investigation, and asocial behaviors were also tallied to produce aggregate scores for each social domain and were subsequently compared between treatment groups.

**Table 1.** Descriptive statistics table for social play. Values for each treatment group are provided as means plus or minus standard error of the mean (SEM).

| Behavior Frequency | AIR | VEH | CAN<sub>THC</sub> | F | df | p | Effect size |
| --- | --- | --- | --- | --- | --- | --- | --- |
| **Play** | 23.4 ± 8.9 | 11.2 ± 9.9 | 14.2 ± 10.6 | 5.830 | 2 | **0.006** | 0.226 |
| Dorsal contact | 5.4 ± 3.7 | 2.6 ± 2.4 | 4.1 ± 3.4 | 2.605 | 2 | 0.086 | 0.115 |
| Evade | 0.9 ± 1.1 | 0.3 ± 0.5 | 1.1 ± 1.5 | 2.641 | 2 | 0.084 | 0.117 |
| Chase | 5.0 ± 3.2 | 1.9 ± 2.5 | 2.2 ± 1.9 | 5.773 | 2 | **0.006** | 0.224 |
| Pin | 2.7 ± 1.5 | 0.9 ± 1.5 | 1.0 ± 1.5 | 7.703 | 2 | **0.001** | 0.278 |
| Pounce | 0.1 ± 0.3 | 0.0 ± 0.0 | 0.4 ± 0.8 | 4.359 | 2 | **0.019** | 0.179 |
| Wrestling | 9.3 ± 5.4 | 5.5 ± 4.8 | 5.4 ± 6.1 | 2.265 | 2 | 0.117 | 0.102 |
| **Social investigation** | 14.4 ± 4.4 | 15.9 ± 3.1 | 14.0 ± 2.5 | 1.140 | 2 | 0.330 | 0.054 |
| Anogenital sniffing | 1.3 ± 1.2 | 1.5 ± 1.2 | 1.5 ± 1.6 | 0.139 | 2 | 0.870 | 0.007 |
| Body sniffing | 13.0 ± 3.8 | 13.7 ± 2.8 | 11.6 ± 2.9 | 1.432 | 2 | 0.251 | 0.067 |
| Social grooming | 0.1 ± 0.4 | 0.7 ± 0.7 | 0.9 ± 1.4 | 2.324 | 2 | 0.111 | 0.104 |
| **Asocial** | 5.5 ± 1.9 | 3.4 ± 2.1 | 4.5 ± 1.7 | 3.812 | 2 | **0.031** | 0.160 |
| Digging | 2.4 ± 2.1 | 1.1 ± 1.1 | 1.6 ± 1.6 | 2.459 | 2 | 0.098 | 0.109 |
| Self grooming | 3.1 ± 1.5 | 2.4 ± 1.5 | 2.9 ± 1.6 | 0.697 | 2 | 0.504 | 0.034 |

#### 2.3.3. Elevated Plus Maze

Behavior in the EPM was assessed during the juvenile period (P27) and during adulthood (P73) using separate cohorts of rats according to previously described methods (Berger et al., 2018; Freels et al., 2020; Walf and Frye, 2007). The EPM apparatus consisted of a raised Plexiglas platform (28.5 inches high) with two open exposed arms and two darker enclosed arms of equal length (21.5 inches/arm; Med Associates Inc., St. Albans, VT). The floors were made of clear Plexiglas and the walls of both closed arms were black. Beginning 2 hr into the dark cycle, rats were transported individually into the testing room and placed at the center of the EPM facing an open arm. The rat was allowed to explore the maze for 5 min while being recorded from above. All tests were run in dim lighting (~10 lux), and behaviors were recorded with Noldus Ethovision XT behavioral tracking software (Noldus, Leesburg, VA). The number of entries and percent time spent in the open and closed arms, and distance traveled in the open arms of the EPM were compared across sexes and treatment groups.

#### 2.3.4. Behavioral Flexibility

An automated attentional set-shifting task was used to assess behavioral flexibility in adulthood. This was conducted using eight 10” × 12” × 12” (L × W × H) Coulbourn Habitest (Holliston, MA) operant chambers each equipped with 2 retractable lever modules, cue lights positioned above each lever, a house light, food trough, and pellet dispenser filled with 45 mg sucrose pellets (Bio-Serv, Flemington, NJ). All rats were single housed beginning on P60 and maintained at 90% free-feeding weight during operant testing in order to incentivize responding for sucrose pellets. Operant training was conducted between P70-110. A small subset of rats from each treatment group (N=25) were injected with latex retrobeads into the basolateral amygdala a minimum of one week prior to the initiation of food restriction, though this had no discernible impact on task performance in these rats. Rats of the same sex were trained and tested in separate male- and female-designated chambers in order to minimize effects of sex-specific olfactory cues.

The attentional set-shifting task was conducted as described previously (Floresco et al., 2008; Brady and Floresco, 2015) (see **Figure 5B** for a schematic of the different training phases). Pre-training began with lever press shaping on a fixed-ratio 1 reinforcement schedule where the left or right lever (counterbalanced between rats) remained extended throughout the session. Rats were required to make a minimum of 60 presses in a single session to reach criterion. After reaching criterion for a given lever, rats were then required to make 60 or more presses on the opposite lever in a single session before advancing to retractable lever training. For retractable lever training, both levers were extended every 20 s, and rats were required to press either lever within 10 s to receive a sucrose pellet. If a rat failed to make a response within 10 s, the levers were retracted and the trial was scored as an omission. All rats received retractable lever training for a minimum of 5 days and continued training until ≤ 5 omissions were committed for 2 consecutive sessions. Rats then performed side preference testing on the same day they completed retractable lever training. Side preference testing consisted of 7 trials (with 2 –8 sub-trials) separated by a 20 s inter-trial interval. During each trial, both levers were extended and a rat’s initial response (left or right lever) was recorded. Side preference was determined by identifying the lever on which the majority of initial responses across the 7 trials were made. Upon completion of pre-training, rats then performed the visual cue discrimination task in which they were required to press the lever associated with an illuminated cue light to obtain the sucrose pellet reinforcer. Rats continued until they made 10 consecutive correct responses. Next, rats performed the set-shifting task that required an extradimensional shift in strategy selection such that they were now required to ignore the visual cue and press the lever opposite to their side preference to obtain the reinforcer. After making 10 consecutive correct responses during set-shifting, rats then advanced to the reversal task. In this task, rats were required to make an intradimensional shift in strategy selection such that they were now required to press the lever opposite to the lever that was reinforced during the set-shifting task to obtain the reinforcer.

For visual-cue discrimination, set-shifting, and reversal tasks, the total number of trials required to reach criterion was tabulated and compared across groups. Error types were also analyzed for set-shifting and reversal tasks by dividing sessions into 16 trial blocks as described previously (Floresco et al., 2008). In the set-shifting task, errors involving the use of the previously reinforced strategy were classified as perseverative errors until fewer than six errors were committed in a block, at which point errors were classified as regressive errors. Never-reinforced errors were classified as when a rat made a response that was not reinforced in either the visual-cue discrimination or set-shifting task. In the reversal task, errors were classified as perseverative until fewer than 10 errors were made in a block. Subsequent errors were then scored as regressive errors. Additionally, it was noted whether perseverative and regressive errors were made toward or away from the cue light during the reversal task.

### 2.4. Statistics

Data were analyzed in SPSS using two-way analyses of variance (ANOVA) with prenatal treatment (CAN_THC_, VEH, AIR) and sex (male, female) as between subject factors. For latency of behaviors in the social play task, we used one-way ANOVAs with treatment as between subject factors due to the number of omissions resulting from frequencies of zero for some behaviors. When significant main effects or interactions were detected, Tukey post-hoc tests were conducted. Alpha was set at .05 for all studies and effect sizes are reported as 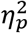.

## 3. RESULTS

### 3.1.1. Maternal vapor exposure affects body weight gain in neonatal offspring

Individual body weights were measured following USV testing on P6, P10, and P13. A two-way ANOVA revealed a main effect of treatment on each day (P6: *F*_(2,76)_ = 9.820, *p* < .001, 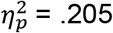; P10: *F*_(2,76)_ = 20.209, *p* < .001, 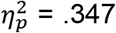; P13: *F*_(2,76)_ = 20.197, *p* < .001, 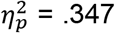; **Fig. 2B**). On P6, AIR pups weighed more than CAN_THC_ and VEH (*p*’s < .05). On both P10 and P13, AIR pups again outweighed CAN_THC_ and VEH neonates, and neonates exposed to CAN_THC_ were also heavier than VEH pups (*p*’s < .05).

**Figure 2.**
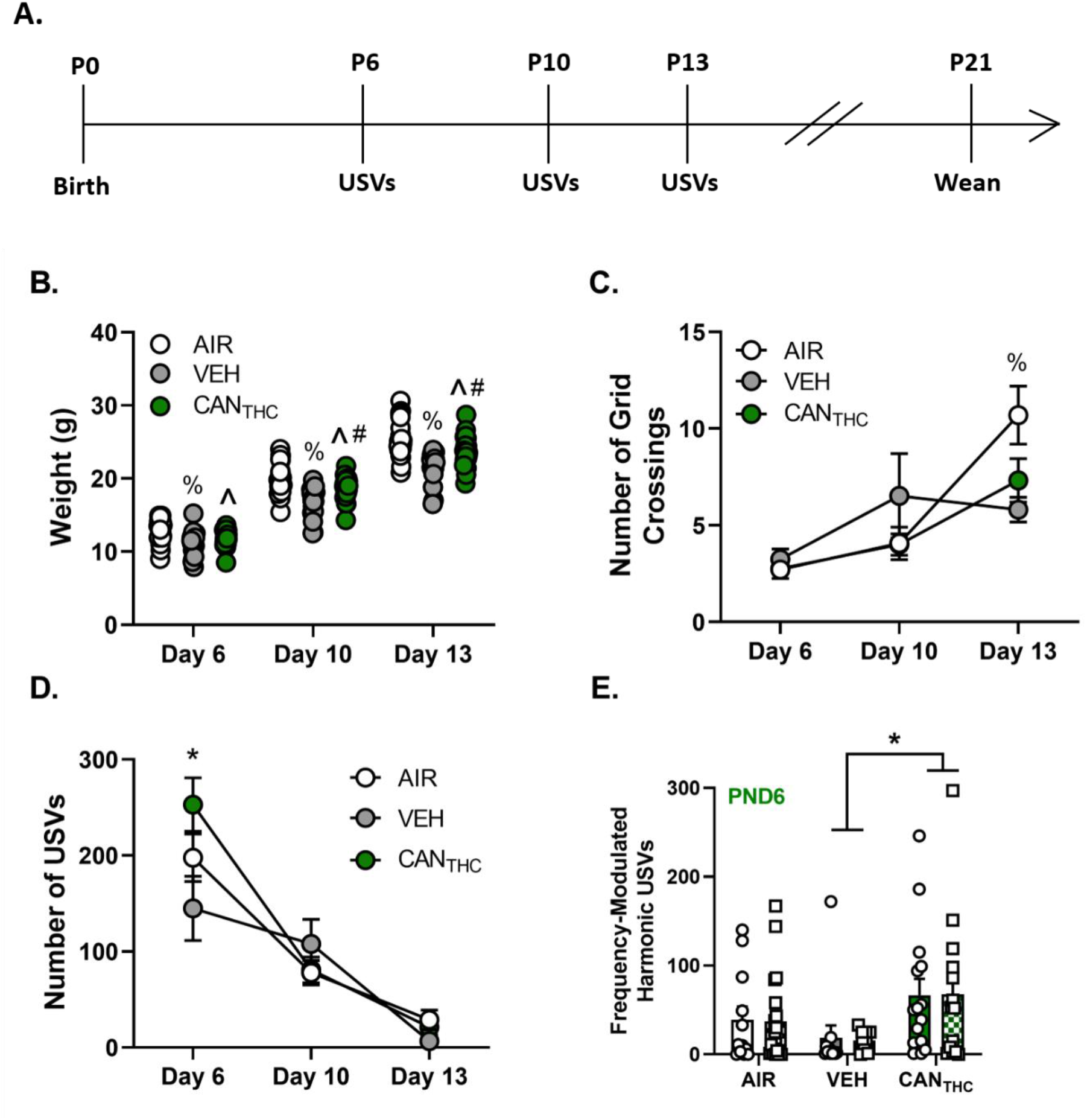
Gestational cannabis exposure leads to altered body weight gain and emotional reactivity in neonate offspring. **(A)** Schematic timeline of neonatal testing timepoints. **(B)** Scatter plot of individual body weights for cannabis vapor- (CAN_THC_; green dots), vehicle vapor- (VEH; gray dots), and no vapor- (AIR; white dots) exposed neonates on postnatal day (P) 6, P10, and P13. **(C)** Mean number of grid lines crossed by CAN_THC_, VEH, and AIR pups during ultrasonic vocalization (USV) testing on P6, P10, and P13. **(D)** Mean number of total USV calls during the 3-min session by CAN_THC_, VEH, and AIR neonates on P6, P10, and P13. **(E)** Mean number of frequency-modulated harmonic calls emitted by CAN_THC_, VEH, and AIR pups on P6. In all graphs, male and female pups are graphed together in their treatment group. * represents a significant difference between the indicated conditions. ^ represents a significant difference between CAN_THC_ and AIR. % represents a significant difference between VEH and AIR. # represents a significant difference between CAN_THC_ and VEH. Data points are presented as mean ± standard error of the mean (SEM).

### 3.1.2. Prenatal vehicle vapor exposure decreases neonatal locomotor activity

The number of grids crossed by pups during USV testing on P6, P10, and P13 was determined. On P13 only, a two-way ANOVA found a main effect of treatment (*F*_(2,76)_ = 3.861, *p* = .025, 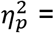 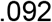; **Fig. 2C**), with neonate VEH pups crossing fewer grids than AIR pups during the USV session on P13 (*p* < .05).

### 3.1.2. Prenatal cannabis exposure increases emotional reactivity during the neonatal period

USVs were recorded on P6, P10, and P13. A two-way ANOVA revealed a main effect of treatment on P6 only (*F*_(2,76)_ = 3.091, *p* = .051, 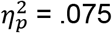, **Fig. 2D**). Pups prenatally exposed to CAN_THC_ made more total calls than VEH (but not AIR) neonates (*p* < .05). Upon further breakdown by number of each call type using a two-way ANOVA, a main effect of treatment was only found for frequency-modulated harmonic calls (*F*_(2,76)_ = 4.615, *p* = .013, 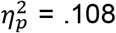, **Fig. 2E**). CAN_THC_ pups emitted more frequency-modulated harmonic calls on P6 than VEH neonates (*p* < .05). No main effects of sex or treatment * sex interactions were observed for any USV measure.

### 3.2.1. Maternal vapor exposure alters social behavior in juvenile male and female offspring

Frequency and latency of play, social investigation, and asocial behaviors during social play sessions were determined. Two-way ANOVA revealed a main effect of treatment on frequency of all play behaviors (*F*(2,40) = 5.830, *p* = .006, 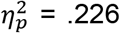; **Fig. 3B**). Specifically, male and female juvenile rats exposed to CAN_THC_ or VEH vapor *in utero* performed fewer total play behaviors compared to AIR controls (*p*’s < .05). This difference may have been driven in part by AIR rats pinning and chasing playmates more than either CAN_THC_ or VEH vapor-exposed rats (pin: main effect of treatment - *F*(2,40) = 7.703, *p* = .001, 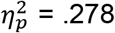 - Tukey’s = *p* < .05; chase: main effect of treatment - *F*(2,40) = 5.773, *p* = .006, 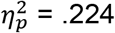 - Tukey’s = *p* < .05; **Table 1**). Additionally, CAN_THC_ juveniles of both sexes pinned playmates more times than did VEH controls (*p* < .05). In addition to decreased frequency of play behaviors, a one-way ANOVA revealed an effect of treatment on latencies to first pin and wrestling (pin: *F*(2,24) = 3.401, *p* = .05, 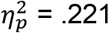; wrestling: *F*(2,35) = 3.991, *p* = .027, 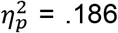; data not shown). Specifically, CAN_THC_ rats initiated their first pin later than AIR controls, and VEH juveniles started wrestling later in play sessions than AIR controls (*p*’s < .05).

**Figure 3.**
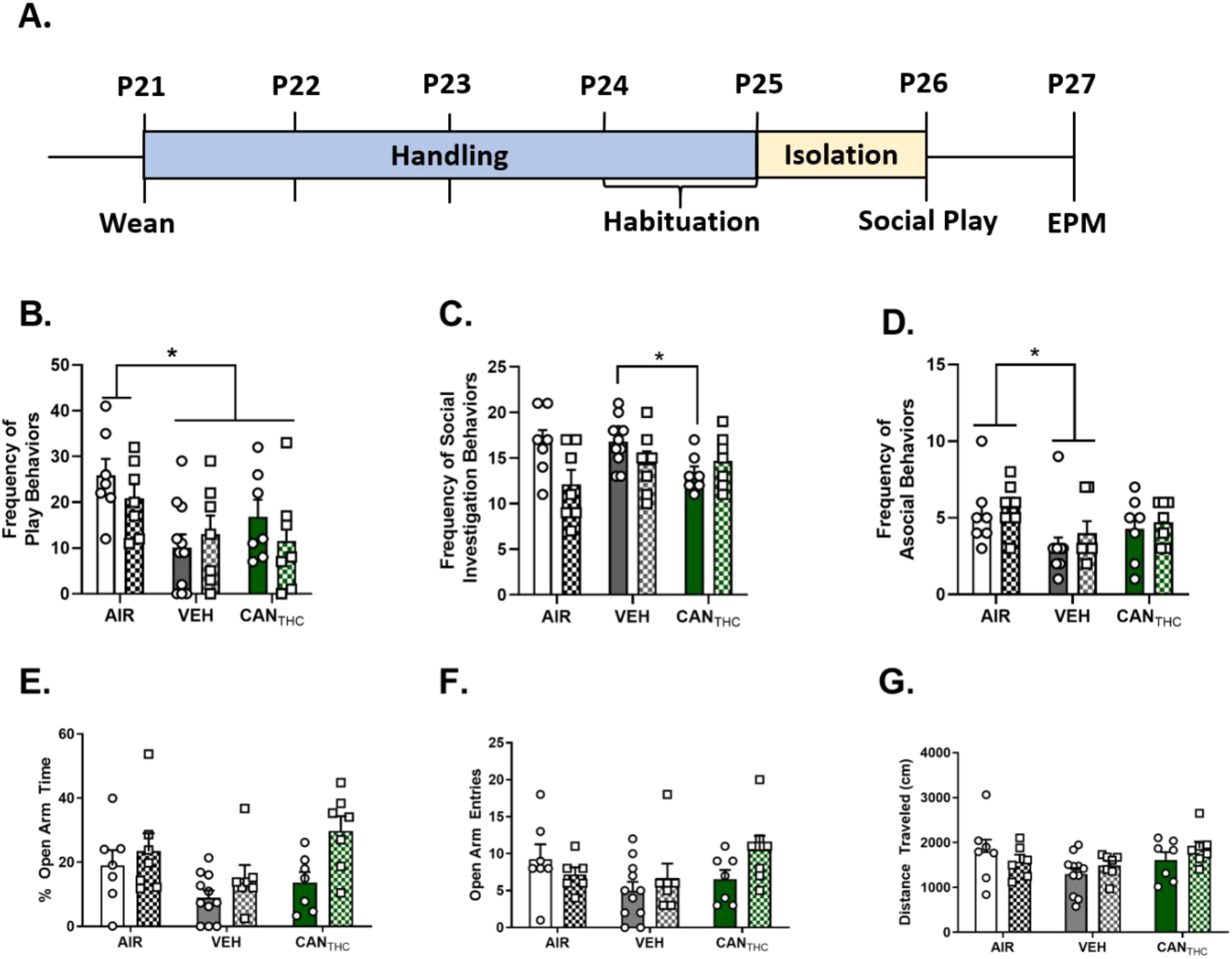
Prenatal cannabis exposure alters social behavior in juvenile rats. **(A)** Schematic timeline of juvenile behavioral testing. Mean frequency of total **(B)** play behaviors (i.e. dorsal contact, pinning, pouncing, following, evading, wrestling), **(C)** social investigation behaviors (i.e., body sniffing, anogenital sniffing, social grooming), and **(D)** asocial behaviors (i.e. digging and self-grooming) exhibited by CAN_THC_, VEH, and AIR exposed rats during the social play session. **(E)** Mean percent of time spent on the open arms of the elevated plus maze (EPM) in CAN_THC_, VEH, and AIR juvenile offspring. **(F)**Mean number of open arm entries made by CAN_THC_, VEH, and AIR juveniles on the EPM. **(G)** Mean distance traveled in cm by CAN_THC_, VEH, and AIR juveniles on the EPM. * represents a significant difference between the indicated treatment groups. On the bar graphs, bars with no patterns represent males and bars with a checkered pattern represent females. NS stands for nonsignificant. Data points are presented as mean ± SEM.

There was a significant treatment * sex interaction for the composite social investigation frequency (*F*(2,40) = 3.301, *p* = .047, 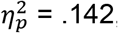, **Fig. 3C**). Male (but not female) CAN_THC_ rats performed fewer total social investigation behaviors during the session compared to male VEH rats (*p* < .05). One-way ANOVA also revealed a main effect of treatment on the latency to first body sniffing (*F*(2,43) = 3.994, *p* = .026, 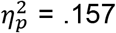, data not shown). Specifically, CAN_THC_ rats of both sexes showed an increased latency to begin body sniffing their playmates (*p* < .05). No significant differences between the total number of individual social investigation behaviors were observed (*p*’s > .05).

Lastly, there was a main effect of treatment on the total number of asocial behaviors (*F*(2,40) = 3.812, *p* = .031, 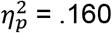; **Fig. 3D**). Specifically, VEH rats of both sexes engaged in fewer asocial behaviors than AIR controls during the play session (*p* < .05). No differences between the individual frequency of digging or self-grooming was noted between any groups (*p*’s > .05).

### 3.2.2. Prenatal vapor exposure does not alter anxiety-like behavior during the juvenile period

Behavior of juvenile rats in the EPM was tracked and used as an index of anxiety-like behavior. Two-way ANOVAs indicated no main effects of treatment, sex, or treatment * sex interactions on percent time exploring the open or closed arms, number of open or closed arm entries, or the total distance traveled in the EPM (all *p*’s > .05, **Fig. 3E-G**).

### 3.3. Prenatal cannabis exposure increases anxiety-like behavior during adulthood

Separate cohorts of male and female rats were tested for anxiety-like behavior in the EPM during adulthood. A two-way ANOVA uncovered a main effect of treatment on the percent of time spent exploring the open arms (*F*(2,46) = 3.843, *p* = .029, 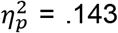, **Fig. 4B**) and the number of open arm entries in adult offspring (*F*(2,46) = 6.738, *p* = .003, 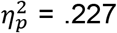, **Fig. 4C**). Specifically, CAN_THC_ rats of both sexes spent less time exploring the open arms of the EPM and made fewer open arm entries than AIR controls (*p*’s ≤ .05). There was no effect of treatment or sex on the total distance traveled in the EPM (*p* > .05).

**Figure 4.**
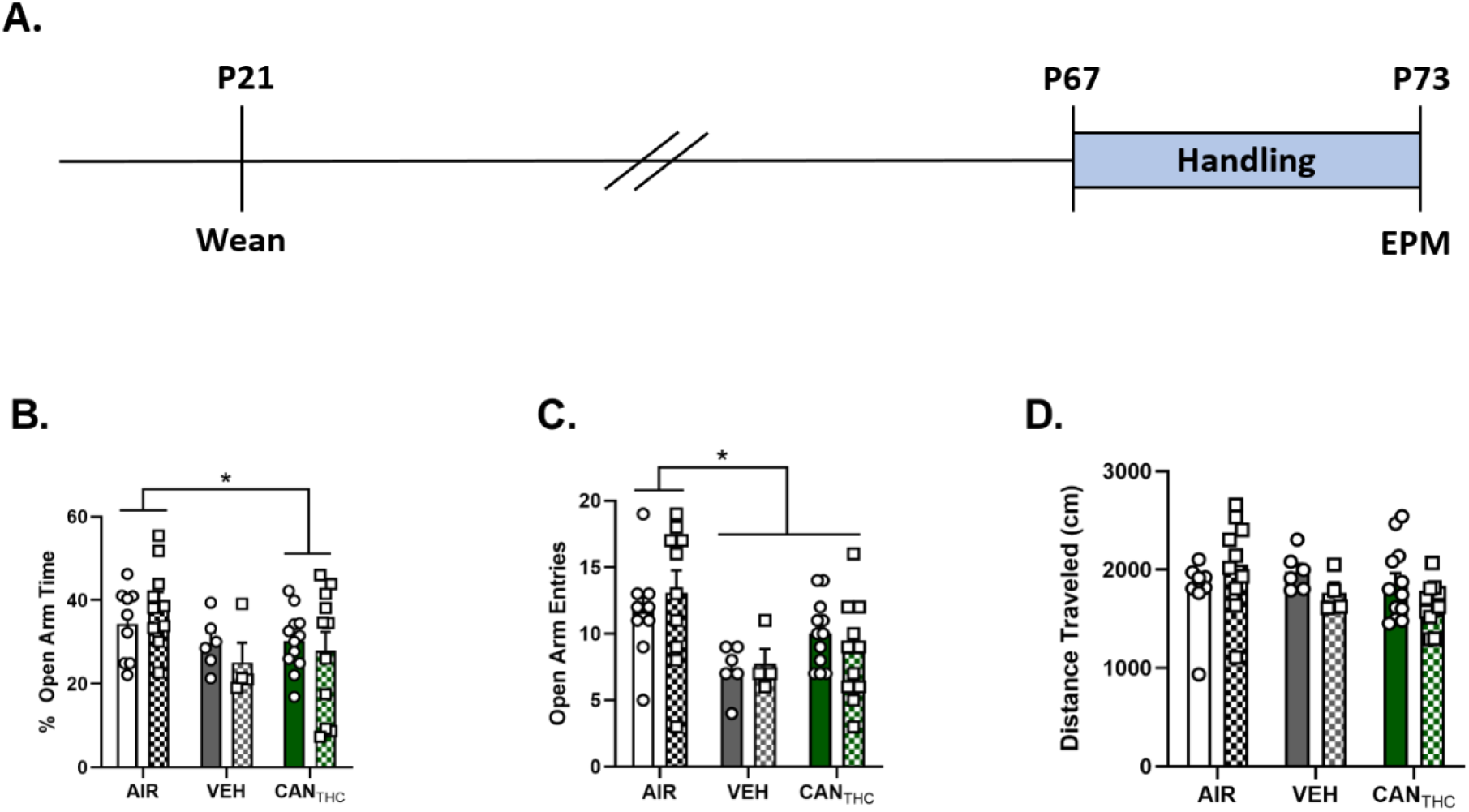
Prenatal cannabis exposure *alters anxiety-like behavior in adulthood.* **(A)** Schematic timeline of adult elevated plus maze (EPM) testing. **(B)** Mean percent of time that CAN_THC_, VEH, and AIR adults spent on the open arms of the EPM. **(C)** Mean number of open arm entries made by CAN_THC_, VEH, and AIR rats on the EPM. **(D)** Mean percent of time that CAN_THC_, VEH, and AIR adults spent on the closed arms of the EPM. **(E)** Mean distance traveled in cm by CAN_THC_, VEH, and AIR adults on the EPM. * represents a significant difference between the indicated treatment groups. On the bar graphs, bars with no patterns represent males and bars with a checkered pattern represent females. Data points are presented as mean ± SEM.

**Figure 5.**
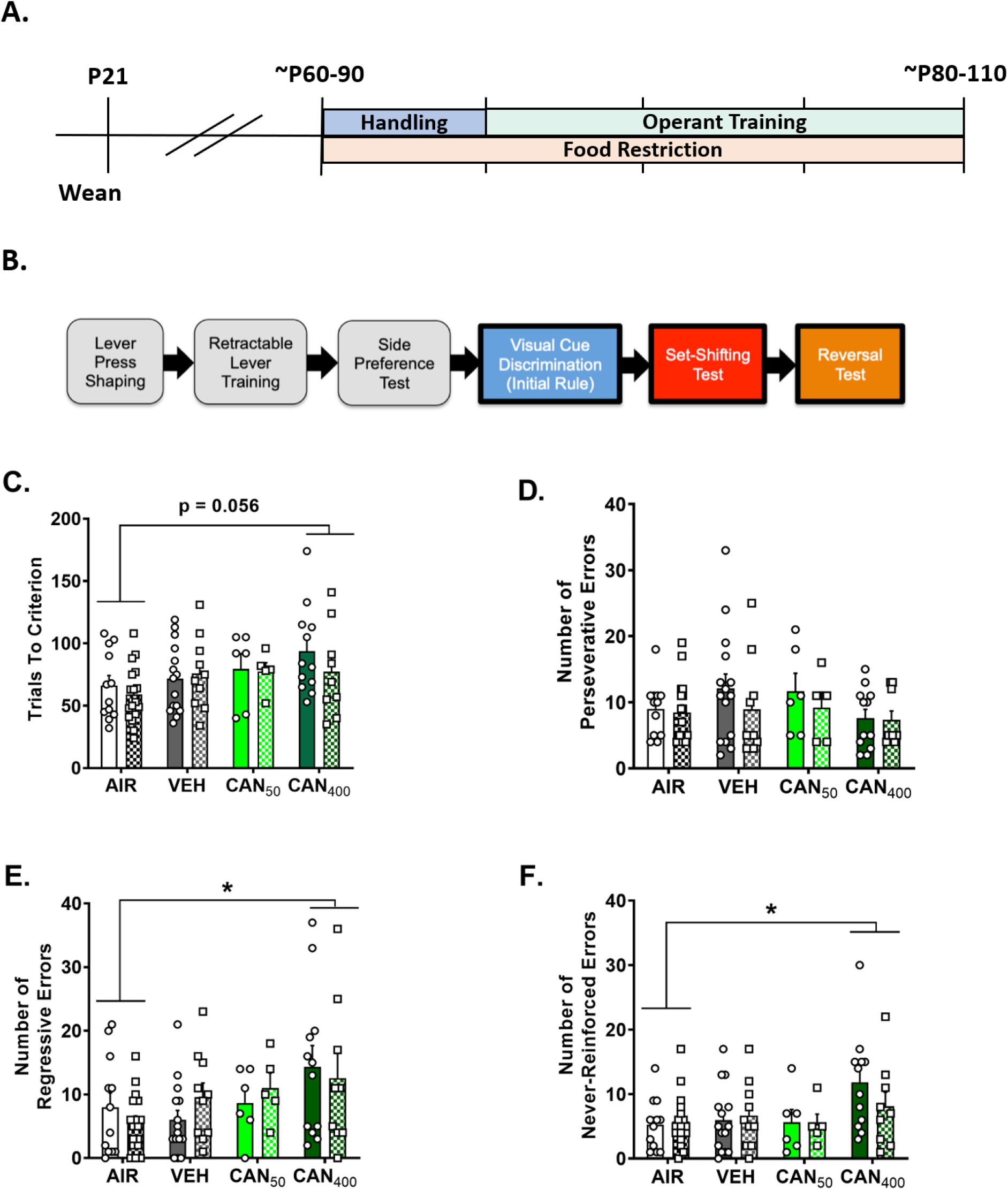
Prenatal cannabis exposure impairs behavioral flexibility in adulthood. **(A)** Schematic timeline of adulthood operant training and behavioral flexibility testing. **(B)** Schematic diagram of the sequence of the different training and testing phases (adapted from Brady and Floresco, 2015). **(C)** Mean number of trials completed by CAN_THC_, VEH, and AIR rats to reach criterion on the set shifting task. **(D)** Mean number of perseverative errors made by CAN_THC_, VEH, and AIR rats on the set shifting task. **(E)** Mean number of regressive errors made by CAN_THC_, VEH, and AIR rats on the set shifting task. **(F)** Mean number of never reinforced errors made by CAN_THC_, VEH, and AIR rats on the set shifting task. * represents a significant difference between the indicated treatment groups. On the bar graphs, bars with no patterns represent males and bars with a checkered pattern represent females. Data points are presented as mean ± SEM.

### 3.4. Prenatal cannabis exposure dose-dependently impairs behavioral flexibility in adulthood

A cohort of rats that did not undergo tests of emotional reactivity or social behavior were tested in the attentional set-shifting task to evaluate effects of prenatal vapor exposure on behavioral flexibility in adulthood. There was no effect of treatment or sex on the acquisition of the initial visual cue discrimination strategy (*p* > .05). However, there was a marginally significant effect of treatment on the number of trials required to meet criterion during the set-shifting task (*F*(3,78) = 2.620, *p* = .056, 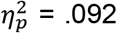, **Fig. 5C**). *A priori* post-hoc analyses comparing task performance in CAN_THC_-exposed rats to VEH and AIR controls revealed that rats exposed to 400 mg/ml (but not 50 mg/ml) CAN_THC_ required significantly more trials to reach criterion than AIR controls (*p* < .05). When error types were examined, two-way ANOVAs revealed a significant main effect of treatment on the number of regressive errors (*F*(3,78) = 3.240, *p* = .026, 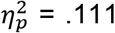, **Fig. 5E**) and never-reinforced errors (*F*(3,78) = 3.290, *p* = .025, 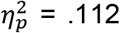, **Fig. 5F**) committed, with CAN_THC_-exposed rats (400 mg/ml) committing more regressive and never-reinforced errors than AIR controls (*p*’s < .05). No main effects of treatment, sex, or treatment * sex interactions were found for perseverative errors (*p* > .05, **Fig. 5D**). There was no effect of treatment or sex on performance in the reversal learning phase or the frequency of error subtypes committed during the reversal task (all *p*’s > .05).

## 4. DISCUSSION

Despite the growing prevalence of cannabis use during pregnancy, relatively little is known about the long-term consequences of prenatal exposure on offspring. In this study, we employed a novel translationally relevant model of maternal cannabis vapor exposure to interrogate the effects of prenatal cannabis exposure on emotional, social, and cognitive endpoints in male and female offspring during early development and into adulthood. Our results indicate that CAN_THC_-exposed offspring made more isolation-induced USVs on P6 relative to VEH-exposed offspring, which is indicative of increased emotional reactivity. Furthermore, we show that male (but not female) CAN_THC_-exposed offspring engaged in fewer social investigation behaviors than VEH-exposed male offspring on P26. In adulthood, CAN_THC_-exposed offspring spent less time exploring the open arms of the EPM and exhibited dose-dependent deficits in attentional set-shifting relative to AIR controls, which indicates long-lasting effects of prenatal cannabis exposure on anxiety-like behavior and behavioral flexibility, respectively. Together, these data support the feasibility of the maternal cannabis vapor exposure model and indicate that this exposure regimen causes enduring effects on the behavioral profile of offspring across the lifespan.

Perhaps one of the most well-established consequences of maternal cannabis use in human populations is lower birth weights in exposed offspring (Gunn et al., 2016). However, animal studies that have used injections of isolated THC or synthetic cannabinoids have failed to observe alterations in neonatal birth weights or body weight gain (Antonelli et al., 2005; Bara et al., 2018; Manduca et al., 2020; Newson & Kelly, 2008; Trezza et al., 2008). Our results indicate that twice daily cannabis vapor exposure during pregnancy leads to decreased body weight gain compared to AIR controls, which is more in line with findings in humans. However, it is important to note that decreased body weight gain was also observed in VEH-exposed offspring, which suggests that this effect may not be caused by cannabis exposure *per se*, but rather by the stress associated with passive vapor delivery. Accordingly, studies in humans have linked prenatal stress to low birth weight (Lima *et al.*, 2018). Surprisingly, CAN_THC_-exposed offspring in our study weighed significantly more than VEH-exposed offspring on P10 and P13, which could indicate that cannabis exposure partially mitigates the effects of vapor-related stress on neonatal body weight gain. These results underscore the need to always include both VEH- and AIR-exposed control groups when employing forced vapor delivery regimens.

In this study, we found that CAN_THC_-exposed neonates emitted more USVs than VEH control pups on P6, which indicates that they are more emotionally reactive to isolation from the nest and littermates. These data are supported by a handful of human studies that have indicated effects of maternal cannabis use during pregnancy on the emotional reactivity of neonate offspring (De Moraes Barros et al., 2006; Fried and Makin, 1987; Lester and Dreher, 1989). While no other study to date has tested cannabinoid-exposed pups at P6, another study conducted by Trezza et al. (2008) similarly documented an increase in the frequency of USVs emitted in THC-exposed offspring at P12. However, other studies have found decreased USVs in pups exposed to the synthetic cannabinoid WIN 55,212-2 (Antonelli et al., 2005; Manduca et al., 2020). It is important to note that, in addition to their use of a synthetic cannabinoid, these studies only recorded calls for the first 15 seconds of isolation in P10 pups, which may partially explain the differences between their findings and ours. Additionally, we found that CAN_THC_-exposed pups emitted more frequency-modulated harmonic calls on P6 than VEH-exposed offspring. Frequency modulation is hypothesized to be a particularly salient characteristic of pup calls that increases the likelihood of being noticed by the dam and assists with pup localization (Brudzynski et al., 1999). Thus, the increased frequency of USVs, particularly of the frequency-modulated harmonic subtype, observed in CAN_THC_-exposed offspring may be advantageous in that it would increase the likelihood that the pup would be retrieved by the dam in the event of separation from the litter. In humans, prenatal cannabis exposure has been linked to increased symptoms of anxiety and depression in preadolescence (Leech et al., 2006). In our study, we did not find any difference between groups for the percentage of time spent on (or the number of entries into) the open and closed arms when juvenile rats were tested in the EPM, which does not support an anxiety-like phenotype. However, our null data are congruent with recent studies in juvenile rats from dams that were subcutaneously injected with THC during gestation (Frau et al., 2019). While these preclinical findings appear to diverge from the human literature, it should be noted that the EPM test only measures a particular facet of unconditioned anxiety-like behavior in rats, specifically that induced by a novel, anxiogenic environment (Walf and Frye, 2013). Thus, the EPM test may not truly capture the domain of anxiety that is experienced in preadolescent human offspring. Unfortunately, to our knowledge, no human studies have examined whether this anxiety phenotype observed in cannabis-exposed offspring extends into adulthood. In contrast to the lack of effect observed during the juvenile period, we unexpectedly found that a separate cohort of male and female rats exposed to CAN_THC_ *in utero* made fewer open arm entries and spent less time exploring the open arms of the EPM compared to AIR controls, which is indicative of increased anxiety-like behavior. Notably, an anxiogenic phenotype has been reported twice before in animal models of prenatal cannabinoid exposure; once using the EPM and another using the open field task (Newson and Kelly, 2008; Trezza et al., 2008). However, two other studies have indicated no differences in EPM behaviors (Bara et al., 2018; Manduca et al., 2020). Again, these incongruent findings may be due to differences in the drug employed, the route of administration, and the maternal exposure regimen.

In contrast to our results from the EPM, juvenile rats exposed to CAN_THC_ *in utero* engaged in less social play behavior and exhibited decreased social investigation, which may be indicative of increased social anxiety during this period. Specifically, CAN_THC_- exposed rats performed fewer play behaviors and pinned playmates later in the session than AIR controls. A decrease in social play behavior has been reported previously in THC-exposed offspring (Trezza et al., 2008), whereas others have reported no differences (Manduca et al., 2020). Accordingly, our data further indicate that male (but not female) offspring exposed to CAN_THC_ engaged in fewer social investigation behaviors than male AIR controls. However, this is somewhat inconsistent with previous rodent studies that have instead found no effects of prenatal cannabinoid exposure on social investigation (Manduca et al., 2020; Trezza et al., 2008). Importantly, the methodology and strategy for quantifying social investigation differed from ours, which may help to explain differences across studies. It is also important to note that social behaviors in VEH-exposed juveniles were also decreased compared to AIR controls in our study. As with neonate body weights, this finding suggests that putative effects of passive vapor exposure also contribute to this social anxiety phenotype in a manner that is independent of the effects of CAN_THC_ exposure. We unexpectedly found that AIR pups engaged in more asocial behaviors (i.e., self-grooming), which would be indicative of a more socially anxious phenotype. However, these bouts of self-grooming typically followed instances of play behavior, which were generally less frequent in vapor-exposed offspring. Therefore, the increase in self-grooming in AIR controls relative to vapor-exposed offspring was likely a byproduct of the increased frequency of play behavior observed in these rats.

In addition to our data revealing long-lasting effects of prenatal cannabis exposure on anxiety-like behavior, we have further shown that these effects also extend to domains of executive functioning. Our results indicate that offspring prenatally exposed to a high dose of CAN_THC_ exhibited impaired set-shifting performance in adulthood relative to AIR-exposed offspring. Importantly, these deficits appear to be selective to flexible responding that requires an extradimensional strategy shift, since no group differences were observed during initial visual cue discrimination learning or the reversal learning components of the task. This behavioral flexibility impairment was primarily due to an increase in the frequency of regressive and never-reinforced errors that were committed. A large body of research has shown that dopaminergic signaling within the corticostriatal circuit is critical for optimal set-shifting performance in behavioral flexibility tasks (Darvas and Palmiter, 2011; Floresco, 2013; Ragozzino, 2002). Accordingly, developmental cannabinoid exposure has been repeatedly shown to alter the ontogeny of dopamine neurons (Bonnin et al., 1994, 1995, 1996; Rodríguez de Fonseca et al., 1991, 1992) and cause augmented dopaminergic activity in humans and laboratory animals (DiNieri et al., 2011; Frau et al., 2019; Rodríguez de Fonesca et al., 1990, 1992). Interestingly, similar increases in regressive and never-reinforced errors have been observed in a set-shifting task following local inactivation of the nucleus accumbens core (Floresco et al., 2006). Moreover, recent studies have indicated that perinatal cannabinoid exposure via maternal injections (Bara et al., 2018) or via breastmilk (Scheyer et al., 2019a) causes significant impairments in endocannabinoid-mediated plasticity in the medial prefrontal cortex (mPFC) of exposed offspring, which likely also gives rise to behavioral flexibility impairments. Thus, the observed deficits in set-shifting performance in the current study are likely attributed to dysfunctional dopaminergic modulation of the corticostriatal circuit that impairs the acquisition and maintenance of newly optimal strategies. Clearly, additional studies are needed to systematically evaluate the effects of prenatal cannabis exposure on corticostriatal function and its precise contribution to the deficits in behavioral flexibility observed herein.

In summary, this study supports the feasibility of using a maternal cannabis vapor delivery regimen to assess emotional, social, and cognitive outcomes across early development and into adulthood. Our findings indicate long-lasting alterations in emotional reactivity, social play behavior, and behavioral flexibility following prenatal cannabis exposure that extend into adulthood. In future studies, it will be important to determine whether these effects are replicated in a volitional cannabis vapor consumption model that effectively eliminates putative effects of stress due to forced drug delivery (Freels et al., 2020). A better understanding of the effects of prenatal cannabis exposure is desperately needed. Our hope is that the knowledge generated from these studies will provide critical information to women and health care professionals about the potential long-term consequences of cannabis use during pregnancy.

## REFERENCES

1. Antonelli, T., Tomasini, M.C., Tattoli, M., Cassano, T., Tanganelli, S., Finetti, S., Mazzoni, E., Trabace, L., Steardo, L., Cuomo, V., Ferraro, L., 2005. Prenatal exposure to the CB1 receptor agonist WIN 55,212-2 causes learning disruption associated with impaired cortical NMDA receptor function and emotional reactivity changes in rat offspring. Cereb. Cortex. 15: 2013–2020. https://doi.org/10.1093/cercor/bhi076.

2. Bailey, J.R., Cunny, H.C., Paule, M.G., Slikker, W. Jr., 1987. Fetal disposition of delta 9-tetrahydrocannabinol (THC) during late pregnancy in the rhesus monkey. Toxicol. Appl. Pharmacol. 90, 315–321. https://doi.org/10.1016/0041-008x(87)90338-3.

3. Bara, A., Manduca, A., Bernabeu, A., Borsoi, M., Serviado, M., Lassalle, O., Murphy, M., Wager-Miller, J., Mackie, K., Pelissier-Alicot, A.L., Trezza, V., Manzoni, O.J., 2018. eLife. 7, e36234. https://doi.org/10.7554/eLife.36234.

4. Berger, A.L., Henricks, A.M., Lugo, J.M., Wright, H.R., Warrick, C.R., Sticht, M.A., Morena, M., Bonilla, I., Laredo, S.A., Craft, R.M., Parsons, L.H., Grandes, P.R., Hillard, C.J., Hill, M.N., McLaughlin, R.J., 2018. The lateral habenula directs coping styles under conditions of stress via recruitment of the endocannabinoid system. Biol. Psychiatry. 84, 611–623. https://doi.org/10.1016/j.biopsych.2018.04.018.

5. Bonnin, A., de Miguel, R., Rodríguez-Manzaneque, J.C., Fernández-Ruiz, J.J., Santos, A., Ramos, J.A., 1994. Changes in tyrosine hydroxylase gene expression in mesencephalic catecholaminergic neurons of immature and adult male rats perinatally exposed to cannabinoids. Brain Res. Dev. Brain Res. 81, 147–150. https://doi.org/10.1016/0165-3806(94)90079-5.

6. Bonnin, A., de Miguel, R., Hernández, M.L., Ramos, J.A., Fernández-Ruiz, J.J., 1995. The prenatal exposure to delta 9-tetrahydrocannabinol affects the gene expression and the activity of tyrosine hydroxylase during early brain development. Life Sci. 56, 2177–2184. https://doi.org/10.1016/0024-3205(95)00205-k.

7. Bonnin, A., de Miguel, R., Castro, J.G., Ramos, J.A., Fernandez-Ruiz, J.J., 1996. Effects of perinatal exposure to delta 9-tetrahydrocannabinol on the fetal and early postnatal development of tyrosine hydroxylase-containing neurons in rat brain. J. Mol. Neurosci. 7, 291–308. https://doi.org/10.1007/BF02737066.

8. Brady, A.M., Floresco, S.B., 2015. Operant procedures for assessing behavioral flexibility in rats. J. Vis. Exp. 96, e52387. https://doi.org/10.3791/52387.

9. Brudzynski, S.M., Kehoe, P., Callahan, M., 1999. Sonographic structure of isolation-induced ultrasonic calls of rat pups. Dev. Psychobiol. 34, 195–204. https://doi.org/10.1002/(SICI)1098-2302(199904)34:3<195::AID-DEV4>3.0.CO;2-S.

10. Campolongo, P., Trezza, V., Ratano, P., Palmery, M., Cuomo, V., 2011. Developmental consequences of perinatal cannabis exposure: Behavioral and neuroendocrine effects in adult rodents. Psychopharmacology. 214, 5–15. https://doi.org/10.1007/s00213-010-1892-x.

11. Coffey, K.R., Marx, R.G., Neumaier, J.F., 2019. DeepSqueak: a deep learning-based system for detection and analysis of ultrasonic vocalizations. Neuropsychopharmacology. 44, 859–868. https://doi.org/10.1038/s41386-018-030306.

12. Darvas, M., & Palmiter, R. D., 2011. Contributions of striatal dopamine signaling to the modulation of cognitive flexibility. Biol. Psychiatry. 69, 704–707. https://doi.org/10.1016/j.biopsych.2010.09.033.

13. De Moraes Barros, M.C., Guinsburg, R., de Araújo Peres, C., Mitsuhiro, S., Chalem, E., Laranjeira, R.R., 2006. Exposure to marijuana during pregnancy alters neurobehavior in the early neonatal period. J. Pediatr. 149, 781–787. https://doi.org/10.1016/j.jpeds.2006.08.046.

14. Dickson, B., Mansfield, C., Guiahi, M., Allshouse, A.A., Borgelt, L.M., Sheeder, J., Silver, R.M., Metz, T.D., 2018. Recommendations from cannabis dispensaries about first-trimester cannabis use. Obstet. Gynecol. 131, 1031–1038. https://doi.org/10.1097/AOG.0000000000002619.

15. DiNieri, J.A., Wang, X., Szutorisz, H., Spano, S.M., Kaur, J., Casaccia, P., Dow-Edwards, D., Hurd, Y.L., 2011. Maternal cannabis use alters ventral striatal dopamine D2 gene regulation in the offspring. Biol Psychiatry. 70, 763–769. https://doi.org/10.1016/j.biopsych.2011.06.027.

16. Floresco, S. B., 2013. Prefrontal dopamine and behavioral flexibility: Shifting from an “inverted-u” toward a family of functions. Front. Neurosci. 7, 1–12. https://doi.org/10.3389/fnins.2013.00062.

17. Floresco, S.B., Block, A.E., Tse, M.T., 2008. Inactivation of the medial prefrontal cortex of the rat impairs strategy set-shifting, but not reversal learning, using a novel, automated procedure. Behav. Brain. Res. 190, 85–96. https://doi.org/10.1016/j.bbr.2008.02.008.

18. Floresco, S.B., Ghods-Sharifi, S., Vexelman, C., Magyar, O., 2006. Dissociable roles for the nucleus accumbens core and shell in regulating set shifting. J. Neurosci. 26, 2449–2457. https://doi.org/10.1523/JNEUROSCI.4431-05.2006.

19. Frau, R., Miczán, V., Traccis, F., Aroni, S., Pongor, C.I., Saba, P., Serra, V., Sagheddu, C., Fanni, S., Congiu, M., Devoto, P., Cheer, J.F., Katona, I., Melis, M., 2019. Prenatal THC exposure produces a hyperdopaminergic phenotype rescued by pregnenolone. Nat. Neurosci. 22, 1975–1985. https://doi.org/10.1038/s41593-019-0512-2.

20. Freels, T.G., Baxter-Potter, L.N., Lugo, J.M., Glodosky, N.C., Wright, H.R., Baglot, S.L., Petrie, G.N., Zhihao, Y., Clowers, B.H., Cuttler, C., Fuchs, R.A., Hill, M.N., McLaughlin, R.J., 2020. Vaporized cannabis extracts have reinforcing properties and support conditioned drug-seeking behavior in rats. J. Neurosci. 40, 1897–1908. https://doi.org/10.1523/JNEUROSCI.2416-19.2020.

21. Fried, P.A, Makin, J.E., 1987. Neonatal behavioural correlates of prenatal exposure to marihuana, cigarettes and alcohol in a low risk population. Neurotoxicol. Teratol. 9, 1–7. https://doi.org/10.1016/0892-0362(87)90062-6.

22. Fried, P.A., Watkinson, B., 1990. 36- and 48-month neurobehavioral follow-up of children prenatally exposed to marijuana, cigarettes, and alcohol. J. Dev. Behav. Pediatr. 11, 49–58. https://doi.org/10.1097/00004703-199004000-00003.

23. Fried, P.A., Watkinson, B., Gray, R., 1998. Differential effects on cognitive functioning in 9-to 12-year olds prenatally exposed to cigarettes and marihuana. Neurotoxicol. Teratol. 20, 293–306. https://doi.org/10.1016/s0892-0362(97)00091-3.

24. Fried, P.A., Watkinson, B., 2001. Differential effects on facets of attention in adolescents prenatally exposed to cigarettes and marihuana. Neurotoxicol. Teratol. 23, 421–430. https://doi.org/10.1016/s0892-0362(01)00160-x.

25. Fried, P.A., Watkinson, B., Gray, R., 2003. Differential effects on cognitive functioning in 13- and 16-year-olds prenatally exposed to cigarettes and marihuana. Neurotoxicol. Teratol. 25, 427–436. https://doi.org/10.1016/s0892-0362(03)00029-1.

26. Gunn, J.K., Rosales, C.B., Center, K.E., Nuñez, A., Gibson, S.J., Christ, C., Ehiri, J.E., 2016. Prenatal exposure to cannabis and maternal and child health outcomes: A systematic review and meta-analysis. BMJ Open. 6, e009986. https://doi.org/10.1136/bmjopen-2015-009986.

27. Hofer, M.A., Shair, H.N., Brunelli, S.A., 2001. Ultrasonic vocalizations in rat and mouse pups. Curr. Protoc. Neurosci. 17, 8.14.1–8.14.16. https://doi.org/10.1002/0471142301.ns0814s17.

28. Holman, P.J., Baglot, S.L., Morgan, E., Weinberg, J., 2019. Effects of prenatal alcohol exposure on social competence: asymmetry in play partner preference among heterogeneous triads of male and female rats. Dev. Psychobiol. 61, 513–524 https://doi.org/10.1002/dev.21842.

29. Javadi-Paydar, M., Nguyen, J.D., Kerr, T.M., Grant, Y., Vandewater, S.A., Cole, M., Taffe, M.A., 2018. Effects of Δ9-THC and cannabidiol vapor inhalation in male and female rats. Psychopharmacology (Berl). 235, 2541–2557. https://doi.org/10.1007/s00213-018-4946-0.

30. Ko, J.Y., Farr, S.L., Tong, V.T., Creanga, A.A., Callaghan, W.M., 2015. Prevalence and patterns of marijuana use among pregnant and nonpregnant women of reproductive age. Am. J. Obstet. Gynecol. 213, 201.e1–201.e10. https://doi.org/10.1016/j.ajog.2015.03.021.

31. Laprairie, R.B., Bagher, A.M., Kelly, M.E., Dupré, D.J., Denovan-Wright, E.M., 2014. Type 1 cannabinoid receptor ligands display functional selectivity in a cell culture model of striatal medium spiny projection neurons. J. Biol. Chem. 289, 24845–24862. https://doi.org/10.1074/jbc.M114.557025.

32. Laprairie, R.B., Bagher, A.M., Kelly, M.E., Denovan-Wright, E.M., 2016. Biased type 1 cannabinoid receptor signaling influences neuronal viability in a cell culture model of Huntington Disease. Mol. Pharmacol. 89, 364–375. https://doi.org/10.1124/mol.115.101980.

33. Leech, S.L., Larkby, C.A., Day, R., Day, N.L., 2006. Predictors and correlates of high levels of depression and anxiety symptoms among children at age 10. Am. Acad. Child Adolesc. Psychiatry. 45, 223–230. https://doi.org/10.1097/01.chi.0000184930.18552.4d.

34. Lester, B.M., Dreher, M., 1989. Effects of marijuana use during pregnancy on newborn cry. Child Dev. 60, 765–771. https://doi.org/10.2307/1131016.

35. Lima, S.A.M., El Dib, R.P., Rodrigues, M.R.K., Ferraz, G.A.R., Molina, A.C., Neto, C.A.P., de Lima, M.A.F., Rudge, M.V.C., 2018. Is the risk of low birth weight or preterm labor greater when maternal stress is experienced during pregnancy? A systematic review and meta-analysis of cohort studies. PLoS One. 13, e0200594. https://doi.org/10.1371/journal.pone.0200594.

36. Manduca, A., Servadio, M., Melancia, F., Schiavi, S., Mazoni, O.J., Trezza, V., 2020. Sex-specific behavioural deficits induced at early life by prenatal exposure to the cannabinoid receptor agonist WIN55, 212-2 depend on mGlu5 receptor signalling. Br. J. Pharmacol. 177, 449–463. https://doi.org/10.1111/bph.14879.

37. McLaughlin, R.J., 2018. Towards a translationally relevant preclinical model of cannabis use. Neuropsychopharmacology. 43, 213–231. https://doi.org//10.1038/npp.2017.191.

38. Metz, T.D., Allshouse, A.A., Hogue, C.J., Goldenberg, R.L., Dudley, D.J., Varner, M.W., Conway, D.L., Saade, G.R., Silver, R.M., 2017. Maternal marijuana use, adverse pregnancy outcomes, and neonatal morbidity. Am. J. Obstet. Gynecol. 217, 478.e1–478.38. https://doi.org/10.1016/j.ajog.2017.05.050.

39. Morales, P., Hurst, D.P., Reggio, P.H., 2017. Molecular targets of the phytocannabinoids: A complex picture. Prog. Chem. Org. Nat. Prod. 103, 103–131. https://doi.org/10.1007/978-3-319-45541-9_4.

40. Navarro, M., Rodríguez de Fonseca, F., Hernández, M.L., Ramos, J.A., Fernández-Ruiz, J.J., 1994. Motor behavior and nigrostriatal dopaminergic activity in adult rats perinatally exposed to cannabinoids. Pharmacol. Biochem. Behav. 47, 47–58. https://doi.org/10.1016/0091-3057(94)90110-4.

41. Navarro, M., de Miguel, R., Rodríguez de Fonseca, F., Ramos, J.A., Fernández-Ruiz, J.J., 1996. Perinatal cannabinoid exposure modifies the sociosexual approach behavior and the mesolimbic dopaminergc activity of adult male rats. Behav. Brain Res. 75, 91–98. https://doi.org/10.1016/0166-4328(96)00176-3.

42. Newson, R.J., Kelly, S.J., 2008. Perinatal delta-9-tetrahydrocannabinol exposure disrupts social and open field behavior in adult male rats. 30, 213–219. https://doi.org/10.1016/j.ntt.2007.12.007.

43. Nguyen, J.D., Aarde, S.M., Vandewater, S.A., Grant, Y., Stouffer, D.G., Parsons, L.H., Cole, M., Taffe, M.A., 2016. Inhaled delivery of Δ^9^-tetrahydrocannabinol (THC) to rats by e-cigarette vapor technology. Neuropharmacology. 109, 112–120. https://doi.org/10.1016/j.neuropharm.2016.05.021.

44. Qato, D.M., Zhang, C., Gandhi, A.B., Simoni-Wastila, L., Coleman-Cowger, V.H., 2020. Co-use of alcohol, tobacco, and licit and illicit controlled substances among pregnant and non-pregnant women in the United States: Findings from 2006 to 2014 National Survey on Drug Use and Health (NSDUH) data. Drug Alcohol Depend. 206: 107729. https://doi.org/10.1016/j.drugalcdep.2019.107729.

45. Ragozzino, M. E., 2002. The effects of dopamine D_1_ receptor blockade in the prelimbic-infralimbic areas on behavioral flexibility. Learn. Mem. 9, 18–28. https://doi.org/10.1101/lm.45802.

46. Richardson, G.A., Ryan, C., Willford, J., Day, N.L., Goldschmidt, L., 2002. Prenatal alcohol and marijuana exposure: Effects on neuropsychological outcomes at 10 years. Neurotoxicol. Teratol. 24, 309–320. https://doi.org/10.1016/s0892-0362(02)00193-9.

47. Roberson, E.K., Patrick, W.K., Hurwitz, H.L., 2014. Marijuana use and maternal experiences of severe nausea during pregnancy in Hawai’i. Hawaii J. Med. Public Health. 73, 283–287.

48. Rodríguez de Fonseca, F., Cebeira, M., Hernández, M.L., Ramos, J.A., Fernández-Ruiz, J.J., 1990. Changes in brain dopaminergic indices induced by perinatal exposure to cannabinoids in rats. Brain Res. Dev. Brain Res. 51, 237–240. https://doi.org/10.1016/0165-3806(90)90280-c.

49. Rodríguez de Fonseca, F., Cebeira, M., Fernández-Ruiz, J.J., Navarro, M., Ramos, J.A., 1991. Effects of pre- and perinatal exposure to hashish extracts on the ontogeny of brain dopaminergic neurons. Neuroscience. 43, 713–723. https://doi.org/10.1016/0306-4522(91)90329-m.

50. Rodríguez de Fonseca, F., Hernández, M.L., de Miguel, R., Fernández-Ruiz, J.J., Ramos, J.A., 1992. Early changes in the development of dopaminergic neurotransmission after maternal exposure to cannabinoids. Pharmacol. Biochem. Behav. 41, 469–474. https://doi.org/10.1016/0091-3057(92)90359-n.

51. Russo, E.B., 2011. Taming THC: potential cannabis synergy and phytocannabinoid-terpenoid entourage effects. Br. J. Pharmacol. 163, 1344–1364. https://doi.org/10.1111/j.1476-5381.2011.01238.x.

52. Scheyer, A.F., Borsoi, M., Wager-Miller, J., Pelissier-Alicot, A.L., Murphy, M.N., Mackie, K., Manzoni, O.J.J., 2019. Cannabinoid exposure via lactation in rats disrupts perinatal programming of the gamma-aminobutyric acid trajectory and select early-life behaviors. Biol. Psychiatry. S0006-3223(19)31662–2.

53. Scheyer, A.F., Melis, M., Trezza, V., Manzoni, O.J., 2019. Consequences of perinatal cannabis exposure. Trends Neurosci. 42, 871–884. https://doi.org/10.1016/j.tins.2019.08.010.

54. Sharapova, S.R., Phillips, E., Sirocco, K., Kaminski, J.W., Leeb, R.T., Rolle, I., 2018. Effects of prenatal marijuana exposure on neuropsychological outcomes in children aged 1-11 years: A systematic review. Paediatr. Perinat. Epidemiol. 32, 512–532. https://doi.org/10.1111/ppe.12505.

55. Smith, A.M., Mioduszewski, O., Hatchard, T., Byron-Alhassan, A., Fall, C., Fried, P.A., 2016. Prenatal marijuana exposure impacts executive functioning into young adulthood: An fMRI study. Neurotoxicol. Teratol. 58, 53–59. https://doi.org/j.ntt.2016.05.010.

56. Spano, M.S., Ellgren, M., Wang, X., Hurd, Y.L., 2007. Prenatal cannabis exposure increases heroin seeking with allostatic changes in limbic enkephalin systems in adulthood. Biol. Psychiatry. 61, 554–563. https://doi.org/10.1016/j.biopsych.2006.03.073.

57. Trezza, V., Campolongo, P., Cassano, T., Macheda, T., Dipasquale, P., Carratù, M.R., Gaetani, S., Cuomo, V., 2008. Effects of perinatal exposure to delta-9-tetrahydrocannabinol on the emotional reactivity of offspring: A longitudinal behavioral study in Wistar rats. Psychopharmacology. 198, 529–537. https://doi.org/10.1007/s00213-008-1162-3.

58. Turner, S.E., Williams, C.M., Iversen, L., Whalley, B.J., 2017. Molecular pharmacology of phytocannabinoids. Prog. Chem. Org. Nat. Prod. 103, 61–101. https://doi.org/10.1007/978-3-319-45541-9_3.

59. Volkow, N.D., Han, B., Compton, W.M., McCance-Katz, E.F., 2019. Self-reported medical and nonmedical cannabis use among pregnant women in the United States. JAMA. 322: 167–169. https://doi.org/10.1001/jama.2019.7982.

60. Walf, A.A., Frye, C.A., 2007. The use of the elevated plus maze as an assay of anxiety-related behavior in rodents. Nat. Protoc. 2, 322–328. https://doi.org/10.1038/nprot.2007.44.

61. Wang, X., Dow-Edwards, D., Anderson, V., Minkoff, H., Hurd, Y.L., 2004. In utero marijuana exposure associated with abnormal amygdala dopamine D2 gene expression in the human fetus. Biol. Psychiatry. 56, 909–915. https://doi.org/10.1016/j.biopsych.2004.10.015.

62. Wen, H., Hockenberry, J.M., Druss, B.G., 2019. The effect of medical marijuana laws on marijuana-related attitudes and perception among US adolescents and young adults. Prev. Sci. 20, 215–223. https://doi.org/10.1007/s11121-018-0903-8.

63. Westfall, R.E., Janssen, P.A., Lucas, P., Capler, R., 2006. Survey of medicinal cannabis use among childbearing women: patterns of its use in pregnancy and retroactive self-assessment of its efficacy against ‘morning sickness.’ Complement. Ther. Clin. Pract. 12, 27–33. https://doi.org/10.1016/j.ctcp.2005.09.006.

